# NIEND: Neuronal Image Enhancement through Noise Disentanglement

**DOI:** 10.1101/2023.10.21.563265

**Authors:** Zuo-Han Zhao, Yufeng Liu

## Abstract

**Motivation:** The full automation of digital neuronal reconstruction from light microscopic images has long been impeded by noisy neuronal images. Previous endeavors to improve image quality can hardly get a good compromise between robustness and computational efficiency.

**Results:** We present the image enhancement pipeline named Neuronal Image Enhancement through Noise Disentanglement (NIEND). Through extensive benchmarking on 863 mouse neuronal images with manually annotated gold standards, NIEND achieves remarkable improvements in image quality such as signal-background contrast (40-fold) and background uniformity (10-fold), compared to raw images. Furthermore, automatic reconstructions on NIEND-enhanced images have shown significant improvements compared to both raw images and images enhanced using other methods. Specifically, the average F1 score of NIEND-enhanced reconstructions is 0.88, surpassing the original 0.78 and the second-ranking method, which achieved 0.84. Up to 52% of reconstructions from NIEND-enhanced images outperform all other 4 methods in F1 scores. In addition, NIEND requires only 1.6 seconds on average for processing 256×256×256-sized images, and images after NIEND attain a substantial average compression rate of 1% by LZMA. NIEND improves image quality and neuron reconstruction, providing potential for significant advancements in automated neuron morphology reconstruction of petascale.

**Availability and Implementation:** The study is conducted based on Vaa3D and Python 3.10. Vaa3D is available on GitHub (https://github.com/Vaa3D). The proposed NIEND method is implemented in Python, and hosted on GitHub along with the testing code and data (https://github.com/zzhmark/NIEND). The raw neuronal images of mouse brains can be found at the BICCN’s Brain Image Library (BIL) (https://www.brainimagelibrary.org).

## 1 Introduction

The advent of high-resolution whole-brain fluorescence microscopy, such as the fluorescence micro-optical sectioning tomography (fMOST) (Li *et al*. 2010; Gong *et al*. 2013, 2016; Zhong *et al*. 2021), has instigated a transformative shift in neuroscience. It empowers researchers with the capability to visualize and scrutinize the neuronal connectome at the mesoscale level (Jiang, Gong and Yuan 2023). However, the immense volume of whole-brain images and the complex nature of axonal arborizations present significant obstacles in capturing neuron morphology (Liu *et al*. 2022). The process of manual reconstruction, a meticulous task of extracting morphology from images, is notably labor-intensive. The potential of automated algorithms to expedite this process is widely acknowledged (Peng *et al*. 2015; Liu *et al*. 2022). Yet, their efficacy remains limited due to their vulnerability to different types of noise and the subpar quality of neuron optical image data (Guo *et al*. 2022).

The noises and artifacts in whole-brain fluorescence microscopy ca‘n be traced back to a variety of factors occurring in image acquisition, including image stitching (Hörl *et al*. 2019), light modulation (Zhong *et al*. 2021), slicing or chemical washing (Zhu *et al*. 2017), and light scattering throughout the brain tissue (Zhu *et al*. 2017). Typical blurs in microscopy (caused by light defocusing and diffraction) can be modeled as the point spread function (PSF), i.e., the impulse response of the imaging system. In particular, whole-brain imaging can introduce an interference arising from mechanical or optical sectioning, whose mechanism is not fully conveyed in current physical PSF models. For example, most versions of fMOST would create an asymmetric flare artifact extending from the excited sample tissues below the knife surface (Zheng *et al*. 2019) (**Figure 1a**). During the image stitching, unnatural boundaries can be left between the large mosaicking artifacts due to uneven illumination (Peng *et al*. 2017b), whose location turns unpredictable in cropped local blocks (**Figure 1b,c**). Auto-fluorescent substances or self-illuminating tissues may also exhibit brightness levels that are on par with neurites in the neuron optical image (**Figure 1d**).

**Figure 1.**
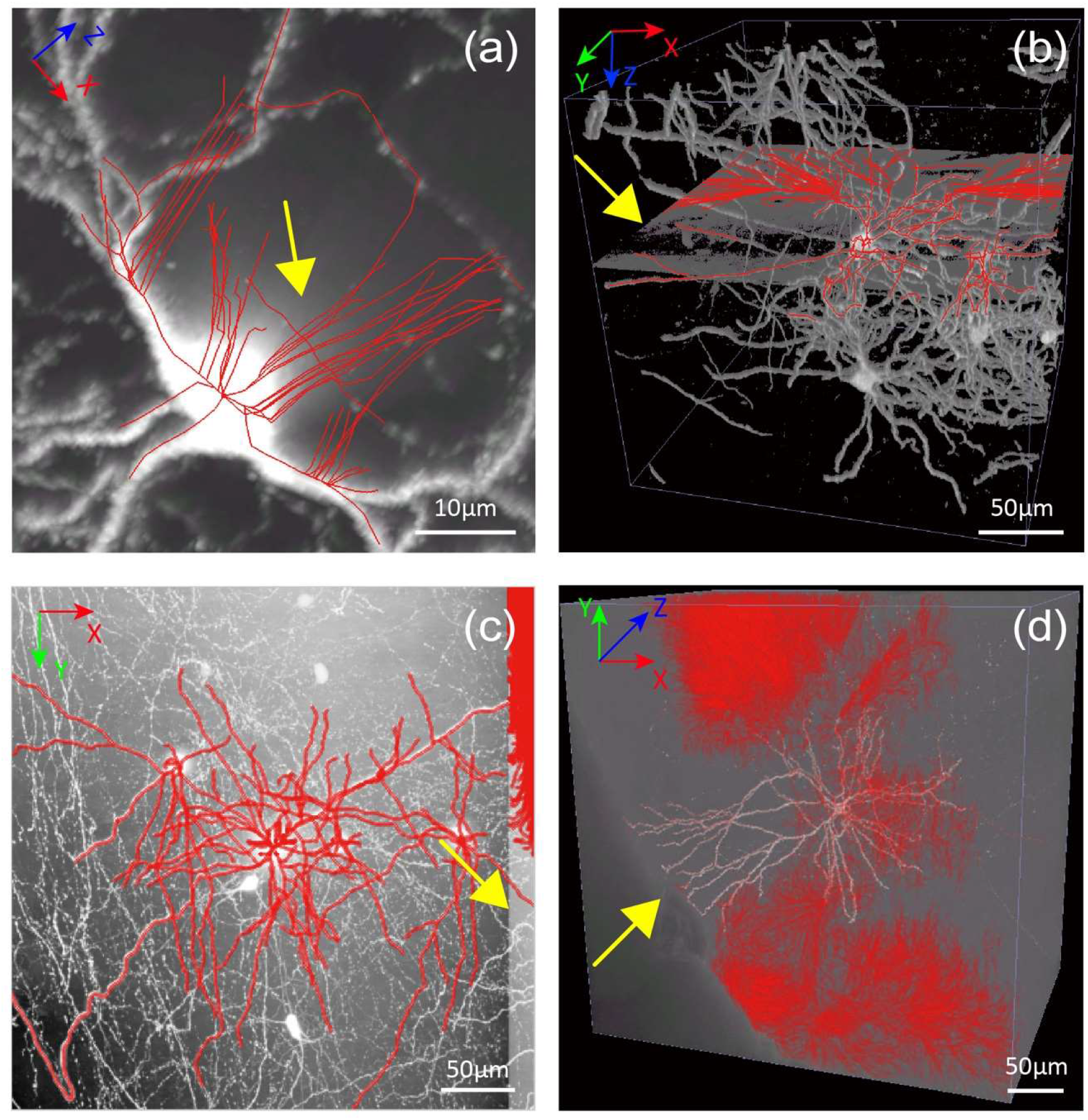
Common noises in light-microscopic neuronal images. Each image is superimposed with auto-reconstruction (utilizing APP2, with default parameters). (**a**) The maximum intensity projection (MIP) of a neuronal image block displaying an attenuated flare artifact along the axial direction-. (**b**) A 3D perspective of an image block with isolated illusion within a plane. Thresholding is applied to the image for better visualization. (**c**) MIP view of a block depicting boundary artifacts between heterogeneous mosaics. (**d**) pervasive background noises. The coordinate system is shown at the top left, and the scale bar at the bottom right. All exemplar images were captured using fMOST techniques, and some noises are specific to fMOST images (e.g., the flare artifact in panel **a**).

Numerous endeavors have been made to reduce noise in neuronal images to enhance tracing quality. To maximally restore the signal, various blind and nonblind deconvolution techniques have been developed (Pankajakshan *et al*. 2009; Ben Hadj *et al*. 2013; He *et al*. 2019). These techniques not only require the accurate measurement or estimation of PSF before or during the deconvolution, but also indicate high computational load. Given the tubular structure of neurites, computational methods have been innovated to locate the foreground directly (Hayman *et al*. 2004; Meijering 2010; Mukherjee and Acton 2015; Zhou *et al*. 2015), but they are computationally intensive as well. Frequency-based filtering can massively save time while achieving less competent enhancement performance (Guo *et al*. 2022). There are also tools tailored for eliminating uneven illumination in bioimages (Smith *et al*. 2015; Chernavskaia *et al*. 2017; Peng *et al*. 2017b), but the mosaicking artifacts can only be handled in a global way. Overall, these existing image processing methods are encumbered by either strict restrictions or high computational cost. With the constant surge in the generation of high-resolution whole-brain images for mammals or primates at the petascale, the urgency for an accurate, robust denoising algorithm that offers affordable computation is escalating.

In this study, such an efficient method is proposed to address the aforementioned issues, named Neuronal Image Enhancement through Noise Disentanglement (NIEND). Results showed that NIEND suppresses the noises and artifacts, and enhances the homogeneity in the foreground, thus facilitating superior performance in automatic tracing. NIEND also saves the computational cost by applying simple but sensitive filters. Its performance has been verified using manually annotated reconstructions and benchmarked against existing methods, showing promise for high-throughput reconstruction of neuron optical images.

## 2 Materials and Methods

### 2.1 Dataset

The study is primarily conducted on the 3D neuron optical image blocks obtained from 39 fMOST brain images (**Supplementary Table S3**, accompanied by 1891 manually annotated reconstructions (Peng *et al*. 2021). The image resolution varies laterally from 0.20 to 0.35μm/pixel while maintaining a consistent resolution of 1μm/pixel in the axial direction. The images were captured in 16-bit. We cropped soma-centered image blocks measuring 1024×1024×256 (voxels, in *XYZ* order) at the highest resolution. This approximates a width ranging from 205μm to 358μm laterally and 256μm axially. These blocks are filtered for sparsity to facilitate the calculation of precision and recall rate of automatic tracing results. It is done by scrutinizing the presence of any significant neurite intersection from other neurons in maximum intensity projection (MIP) of the XY-plane. A total of 863 neurons was left after the selection (**Supplementary Table S3**). Manual reconstructions (gold standards) were cropped accordingly, with any suspending structures (those with parent structures outside the block) removed. To validate the effectiveness of NIEND on other image modalities, we also performed benchmarks on the BigNeuron dataset (Peng *et al*. 2015) with 162 microscopy images of various image types, including confocal, two-photon and epi-fluorescent imaging.

### 2.2 Overview of NIEND Pipeline

The NIEND pipeline is composed of three key modules: high-pass filtering, intensity shifting, and low-pass filtering (**Figure 2**). The high-pass filtering module incorporates multiple filters to combat various types of low-frequency noise. In this study, we showcased two filters—the diffusion filter and the orthogonal filter—to respectively eliminate the attenuation noise (**Figure 1a**, consisting of the PSF response and the flare artifact, extending towards the negative direction of the Z-axis, i.e., the opposite direction of sample sectioning) and the orthogonal artifact (**Figure 1b–d**, large, homogeneous or orthogonally situated artifacts). Afterwards, neuronal images may still suffer from an intense illumination heterogeneity, resulting in the loss of weak neurites during bit depth downgrade (typically 16-bit to 8-bit) and auto-reconstruction. Therefore, the intensity shifting module is employed to homogenize the uneven foreground and introduces a instance-aware mechanism for enhancing very weak neurons. Facilitated by the narrowed dynamic range, a near-lossless bit depth downgrade to 8-bit is performed subsequently. Finally, the low-pass filtering module uses a wavelet filter to mitigate remaining high-frequency noise.

**Figure 2.**
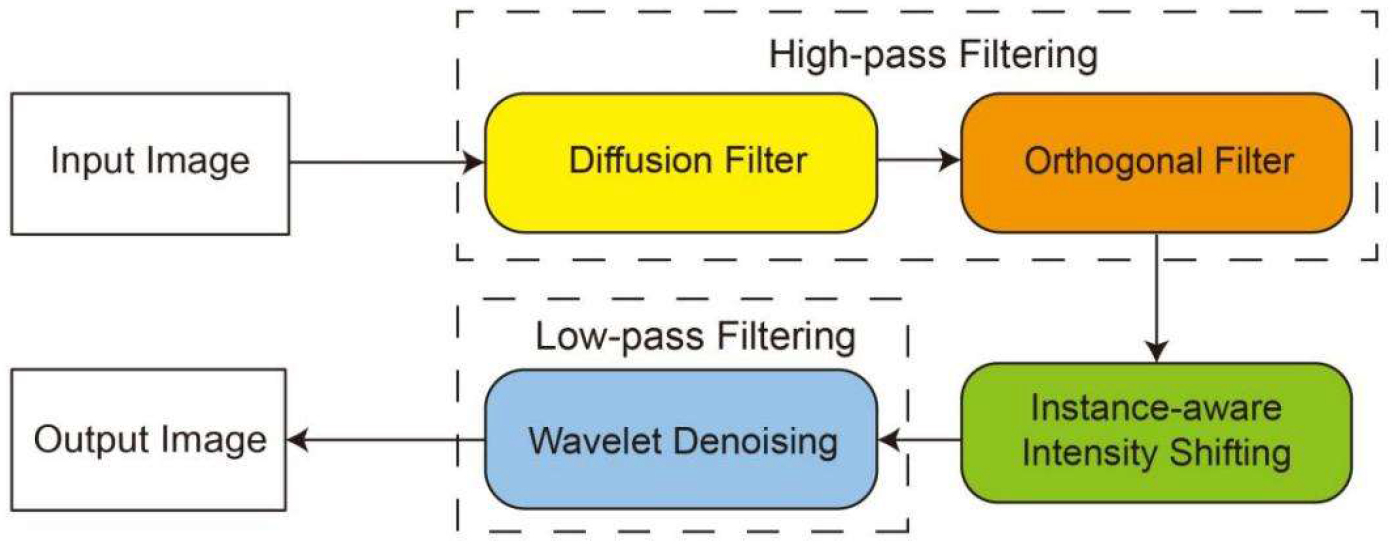
Schematic pipeline of NIEND. This diagram presents a simplified flow of the NIEND, starting from the input image. At the start is a high-pass filtering module integrating two filters: the diffusion filter and the orthogonal filter. These filters are engineered to respectively eliminate the attenuation noise (consisting of the PSF response and the flare artifact) and orthogonal artifacts (large, homogeneous or orthogonally situated artifacts). Subsequent steps include instance-aware intensity shifting and wavelet denoising, which refine the high-pass filtered result by adjusting the dynamical range and eliminating potential high-frequency noise.

### 2.3 sNoise-specific high-pass filtering

The high-pass filtering module is a critical step to refine the image quality. It starts with a diffusion filter (**Figure 3)**. This filter restores the neurite signal by iteratively removing the attenuation noises from each Z-slice (**Equation 1.1.1**). For the n-th slice, the input is denoted as **raw**_**n**_, the attenuation noise as **noise**_n_ and the output as **res**_**n**_. Considering the asymmetry of the attenuation noise, we define **noise**_**n**_ as the degradation of all the previous restoration results (**Equation 1.1.2**). To disentangle the degradation on each slice, the restoration is transformed into **Equation 1.1.3** by assuming that the degradation is additive and escalates only with respect to the axial distance to the current slice (**dist**_**i**_).

**Figure 3.**
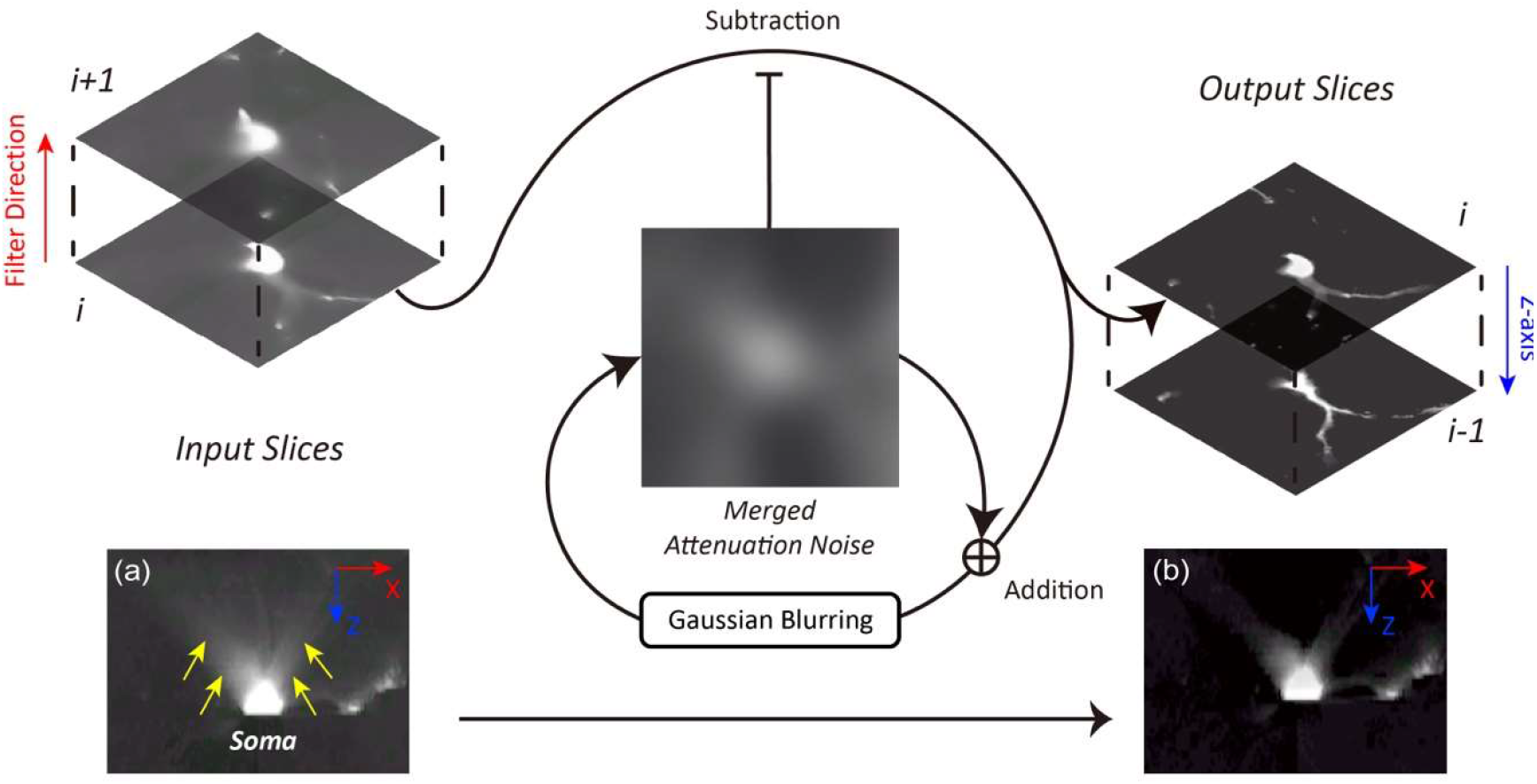
Schematic steps of the diffusion filter. The filtering sequentially subtracts an updating slice of attenuation noise from each raw image slice along its Z-axis, typically the sectioning direction. The circulation denotes that the attenuation noise is recursively updated during each iteration, by merging with the denoised output of this iteration and implementing Gaussian blurring. (**a**) an XZ-slice of an input image, showing a flare artifact extending along the Z-axis and dispersing over the X-axis (yellow arrows). (**b**) the processed image where the artifact is significantly reduced.

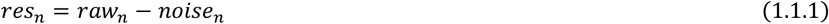

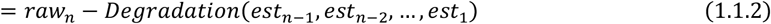

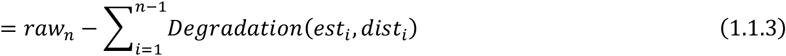

Then, we approximate the degradation as thermal diffusion, by introducing a Gaussian blur (**Equation 1.2.1**) with its standard deviation (σ) escalating with **dist**_**i**_. As the slices are uniformly spaced, σis transformed as (**n** – **1**)**s**, where **s** is a tunable parameter for the diffusion speed. In this study, **s** is determined as inversely proportional to the lateral resolution of the image (**Supplementary Table S1**). Given its associativity property, we can conveniently use the same Gaussian kernel to compose all the degradation processes (**Equation 1.2.2**).

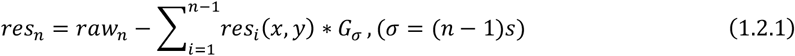

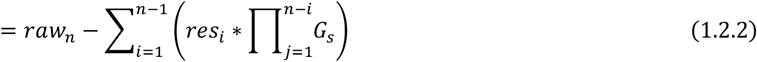

The Gaussian kernel offers another benefit of reducing the computational cost by facilitating a recursion definition (**Equation 1.3.1 through 1.3.3**).

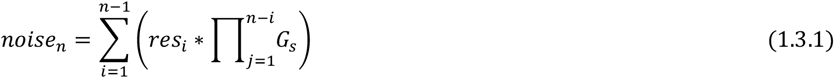

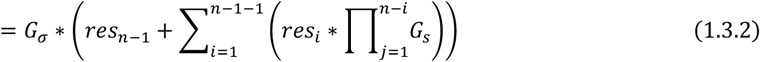

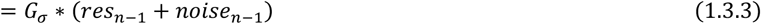

Now, the restoration of each slice can be performed only incorporating the immediate predeceasing result and an updating slice of the attenuation noise (**Equations 1.4**). For the adaptiveness across the slices, we add a multiplier α(**n**) to dynamically match the average intensity of **noise**_**n**_ with that of the current slice. This multiplier is allowed to be tuned with a parameter, **k** (set as 0.9 in this study), so as to neutralize the filter performance (e.g. **k** as 1.0 brings superb but disruptive denoising effect), in consideration that the tracing algorithm in use might be sensitive to signal breakups.

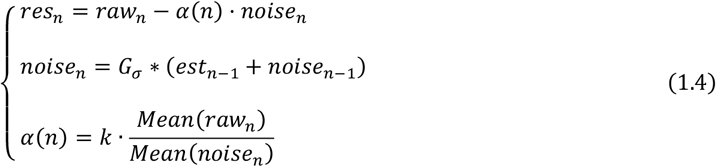

Subsequently, an orthogonal filter is applie‘d. It acquires two average profiles on the YZ-plane and the XZ-plane, which are then combined into a 3D array of the original image shape through the direct sum operation. By subtracting this array, the image becomes free of low-frequency stripes (**Supplementary Figure S1**), large homogeneous region (**Supplementary Figure S4a**), clear boundary artifacts (**Supplementary Figure S4c**), etc. The schematics and results of each step are also depicted in **Supplementary Figure S1**.

### 2.4 Instance-aware intensity shifting

The instance-aware intensity shifting is performed to determine the optimal dynamic range for the image, enhance the foreground homogeneity and lower the image size. The lower bound is set as a percentile that clears most of the background volume, given the sparsity of neurites. Empirically, a percentile around 1.0% is practical for a dendrite block of sub-micrometer resolution. In this study, minor adjustments (±0.5%) are made for images cropped from brains of different resolutions (**Supplementary Table S1**). The upper bound should be sufficiently small to maximally retain the details in the final 8-bit conversion as well as to enhance the signal for very weak neurons. Therefore, an instance-aware mechanism is introduced by taking it as the lesser between 255 above the lower bound and half the maximum intensity of the soma region, which we take as an image block of size 128×128×32 voxels, equivalent to a real size of 25.6×25.6×32–44.8×44.8×32μm^3^. However, for some tracing algorithms (e.g., APP2), this mechanism is actually unnecessary, and the 255 above the lower bound is enough. Finally, the intensity of the whole image will be clipped and linearly rescaled between the bounds.

### 2.5 Wavelet low-pass filtering

High-frequency noises, including the irregular and fractured residual artifact that unaddressed in previous steps, can be staged during the intensity shifting as a side effect (**Supplementary Figure S2a**). Here, wavelet denoising (Halidou *et al*. 2023) is utilized for its little to none degradation to neuronal structures compared with other typical low-pass filters. It can also fix glitches (1–2 voxels of abrupt change in intensity), including minor attenuation noise and breakups (**Supplementary Figure S2c**). The denoising is implemented as slice-wise to minimize memory usage. In detail, we adopt the Haar wavelet and hard thresholding. The wavelet level is capped at 2 to model the resolution difference. The thresholds for each wavelet level are adaptively estimated by BayesShrink (Chang, Bin Yu and Vetterli 2000).

### 2.6 Benchmarking

NIEND is tested on 863 sparsely labeled mouse neuronal blocks and 162 BigNeuron images, in comparison with various existing techniques, including adaptive thresholding (which subtracts the average intensity of neighboring voxels), multiscale enhancement (Zhou *et al*. 2015), and Guo’s enhancement method (Guo *et al*. 2022). The reconstruction of neurons is conducted through all-path-pruning 2 (APP2) (Xiao and Peng 2013), with the default parameter settings. Additional comparisons with the PSF-based Richardson-Lucy (R-L) deconvolution and benchmarks on other tracing algorithms, including APP1 (Peng, Long and Myers 2011) and NeuroGPS-Tree (Quan *et al*. 2016), are also performed. The protocols for the additional experiments are given in **Supplementary Methods**.

To compare the image quality, we computed the signal-background contrast (SBC) and the within-image homogeneity (WIH) as presented in previous studies (Guo *et al*. 2022). The foreground is estimated as a neuronal mask derived from the radius-profiled manual reconstruction, while the background is defined as the difference between the two-fold-enlarged neuronal mask and the foreground. Based on these masked voxels, the SBC is calculated as the ratio between the median of the foreground and the median of the background (**Equation 2.1**). The background WIH is calculated as the uniformity (**Equation 2.2**) of the normalized histogram of the background intensities (after z-normalizing the whole image), while the foreground WIH is calculated as the relative standard deviation (RSD) of the foreground intensities (**Equation 2.3**). Some metrics have been modified to avoid zero division.

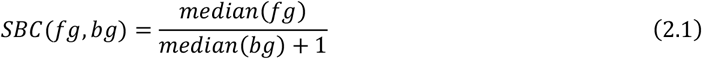

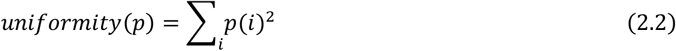

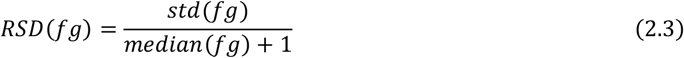

The precision, recall and F1 are computed for every neuronal block. These calculations are based on the Percent of Different Structures (PDS) metric (Peng, Long and Myers 2011; Xiao and Peng 2013), a rate measuring the percentage of deviations of a reconstructed neuron against a reference neuron (**Equation 2.4**). The deviation is quantified as the number of the components whose distances to the nearest reference point exceeds a limit (**Equation 2.5**). In this study, the limit is specified as 15 voxels (approximately 3μm).

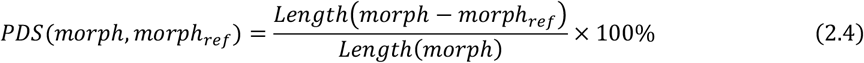

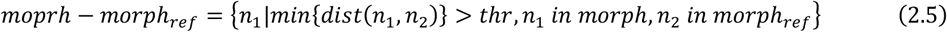

Now, the precision is determined from the error, which is the PDS of the reconstruction against the gold standard (**Equation 2.6**) while the recall is determined by reversing the gold standard and the reconstruction (**Equation 2.7**). F1 is computed accordingly (**Equation 2.8**).

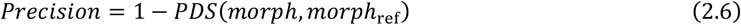

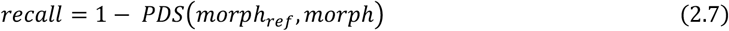

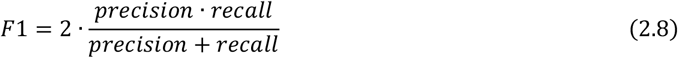

In this assessment, the optimal image depths are adopted in automatic tracing for each technique (estimated by comparing the tracing metrics). To be specific, the 8-bit version is used for multiscale enhancement and Guo’s method, while the 16-bit version is used for the remaining methods. Note that APP2 automatically converts any image to 8-bit before tracing. Running time and memory usage tests are conducted on 30 randomly sampled image blocks serially, with the first 10 results discarded to prevent initialization bias. The running time and memory usage are also acquired for image blocks of a more regular size (256×256×256 voxel^3^).

## 3 Results

### 3.1 NIEND improves the image quality substantially

We showcased the outputs of different NIEND steps with examples containing typical noises and artifacts discussed (**Figure 4**). The application of high-pass filtering significantly reduced the image noise and artifacts while retaining the neurite signals (**Figure 4b**). After the intensity shifting, the neurite structures are accentuated (**Figure 4c**), albeit with an increase in some of the remaining noises, which are then mitigated with wavelet denoising (**Figure 4d**).

**Figure 4.**
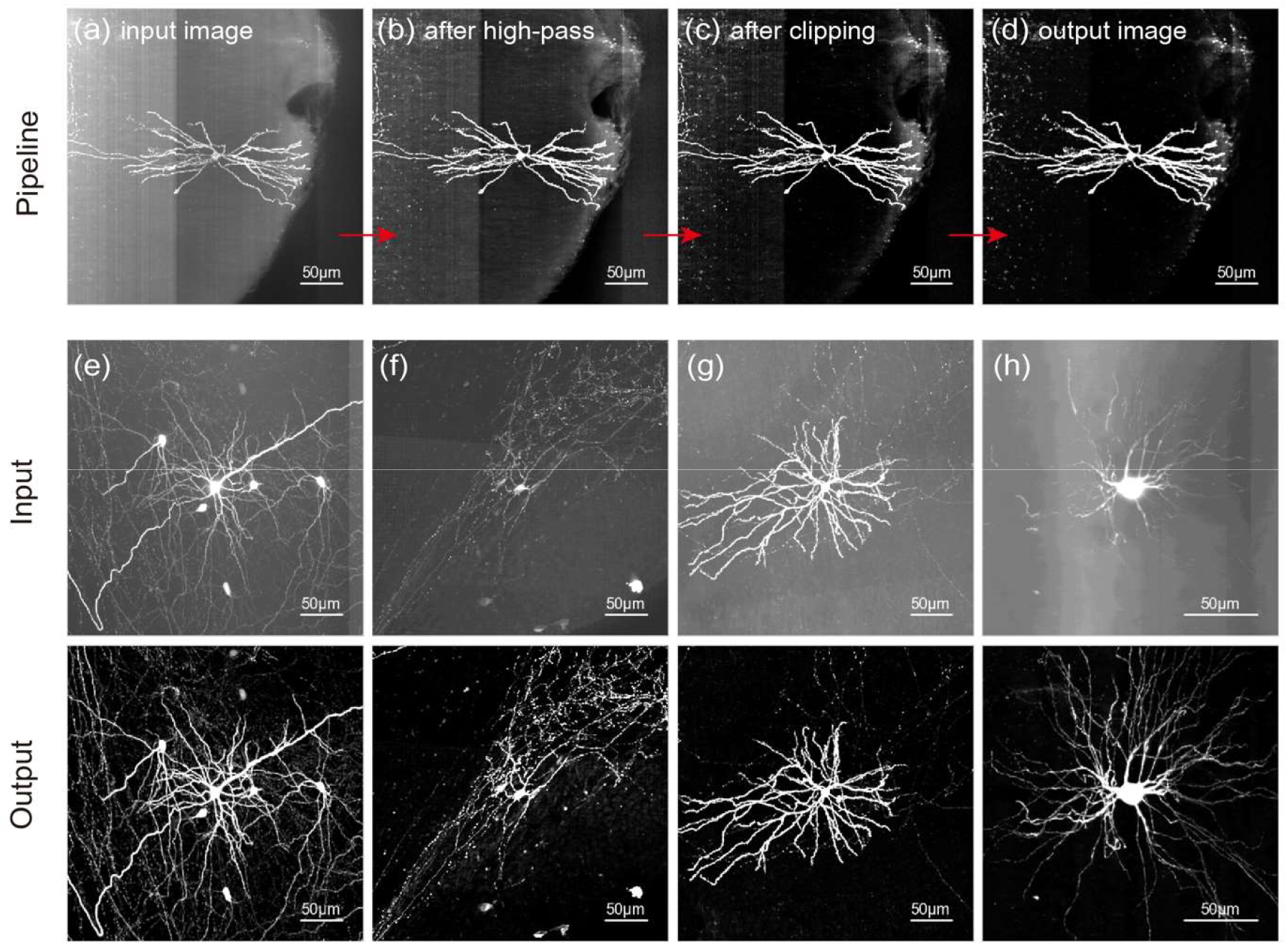
Examples of images after NIEND. (**a–b**) An example in MIP that illustrates the output at each stage of the proposed pipeline. This encompasses an initial input image (**a**), the image after high-pass filtering (**b**), the image after instance-aware intensity shifting (**c**), and the final output image (**d**). (**e–h**) Comparison of another four initial (input, top) and enhanced images (output, down) pairs using NIEND. All images are z-score normalized and converted to the unsigned 8-bit range (0 - 255).

More visual examples confirm the improvement of NIEND (**Figure 4e–h**). It eradicates substantial quantities of non-neuron artifacts, such as the illuminated possible brain tissues (**Figure 4e,f**) and the sinusoidal band (**Figure 4g,h**). Several other types of issues that pose challenges for tracing are also addressed, such as poor contrast between the cells and background (**Figure 4e**), thin and dim local axon (**Figure 4f**), and broad dispersion between the foreground structures (**Figure 4g,h**).

### 3.2 Comparison with other image enhancements

We further compared the performance of NIEND with existing state-of-the-art methods, including adaptive thresholding (AdaThr), multiscale enhancement (Multiscale), and Guo’s enhancement method (Guo). The classic methods like the anisotropic filter and the Meijering filter, despite their power, are excluded for their extreme computational cost.

The advantages of NIEND are demonstrated with seven neuronal image blocks with representative noise types (**Figure 5**). Adaptive thresholding is a highly lightweight form of high-pass filtering. It excels at denoising while introducing side effects, including hollow soma and disruption of nearby stems (**Supplementary Figure S3a**), because it can only perform very specific elimination of a fixed frequency range that often includes neurite features. Its disruptive issue even escalates when the image quality is high. Its denoising also overlooks the boundary artifact (**Supplementary Figure S3d**). In contrast, NIEND is capable of settling these issues with an improved denoising performance (**Supplementary Figure S3c**,**f**). Multiscale enhancement effectively denoises the samples (**Figure 5a**) but results in a catastrophic loss when it misidentifies neurites as noise (**Figure 5d–f**).

**Figure 5.**
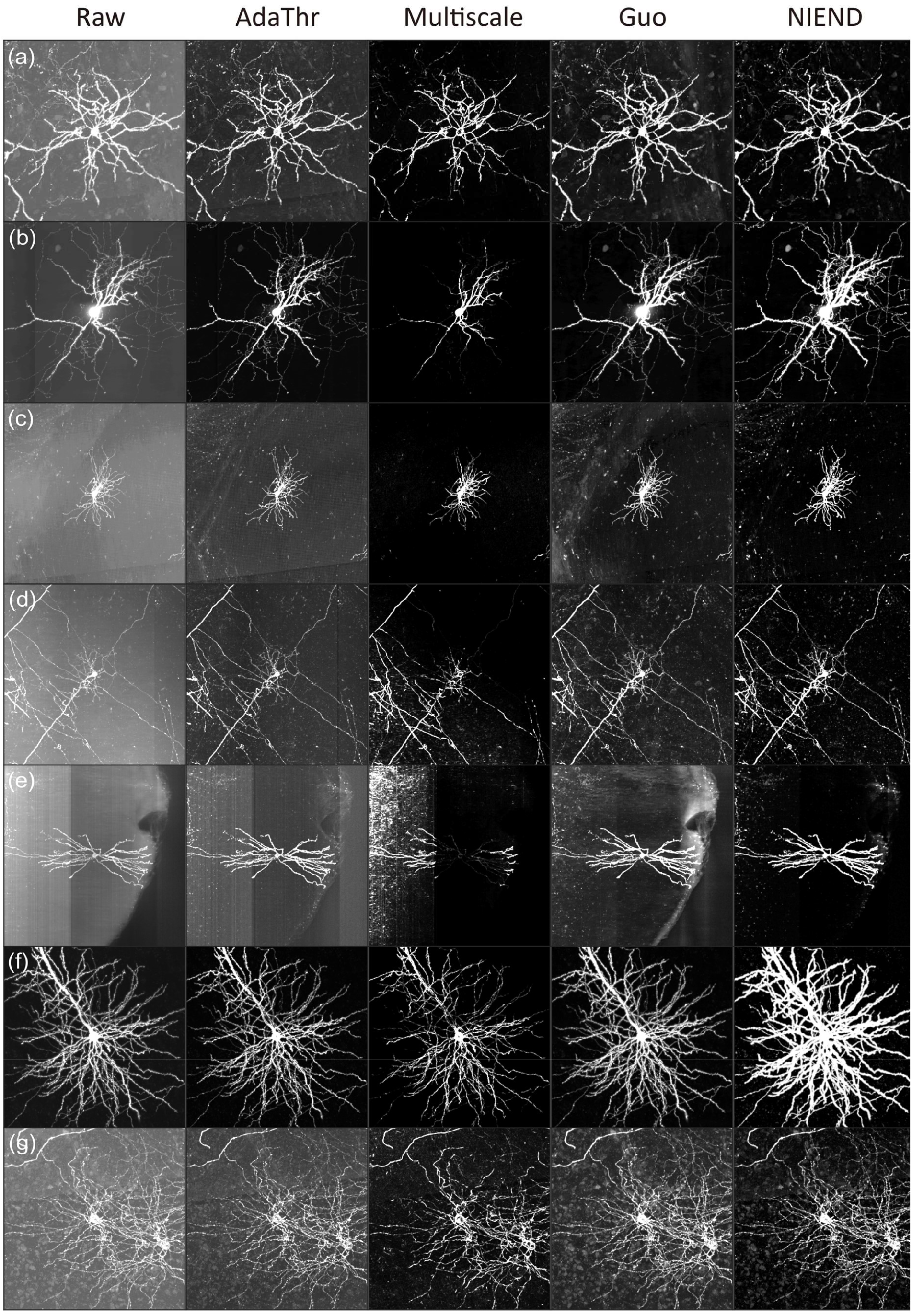
Comparison between NIEND and existing algorithms. Each column represents one method, while each row represents an image. All images are z-score normalized and converted to unsigned 8-bit range (0 - 255). From left to right, the raw images, images after adaptive thresholding, images after multiscale enhancement, images after Guo’s method, and images after NIEND. Each sample represents a typical combination of noises: (**a**) high background noise. (**b**) a high-intensity soma and faint axons. (**c**) high, uneven background noise. (**d**) weak neurites while strong and uneven background noise. (**e**) noisy background with orthogonal and irregular boundaries. (**f**) low noise level. (**g**) interwound neurons with high level of noises.

Guo’s method is a competent tool for weak neurite enhancement. Still, its efficacy is comparatively limited in terms of background noise removal. Guo’s method claims to denoise the slowly varying background noise with high-pass filtering in Fourier space, but omits much of the irregular noise (**Figure 5c–e**). Instead, NIEND demonstrates its advantage in enhancing the weak signal, including weak axons (**Figure 5b**) and neurites obscured by strong background noise (**Figure 5d**). The intensity among neurites is homogenized as well (**Figure 5f**).

The image quality is also compared quantitatively (**Figure 6**). NIEND exhibits the largestincrease in the signal-background contrast (SBC), with 40-fold larger than the raw image and 14% higher than the second-ranking method (adaptive thresholding). The metric SBC is analogous to the Signal-to-Noise Ratio (SNR) and was utilized as a primary metric for calibrating neuronal image enhancement in previous studies (e.g., Guo *et al*. 2022). In terms of background noise reduction, NIEND secures the second-highest gain in background uniformity (10-fold larger than that of raw image). Regarding the foreground relative standard deviation (RSD), whose increase indicates the rise of signal loss, all of the methods display degradation to some degree. NIEND achieves minimal signal loss with only a slightly larger value (2.1) than that of traced morphologies using raw images (1.5). Overall, NIEND delivers the most significant and lossless enhancement in image quality.

**Figure 6.**
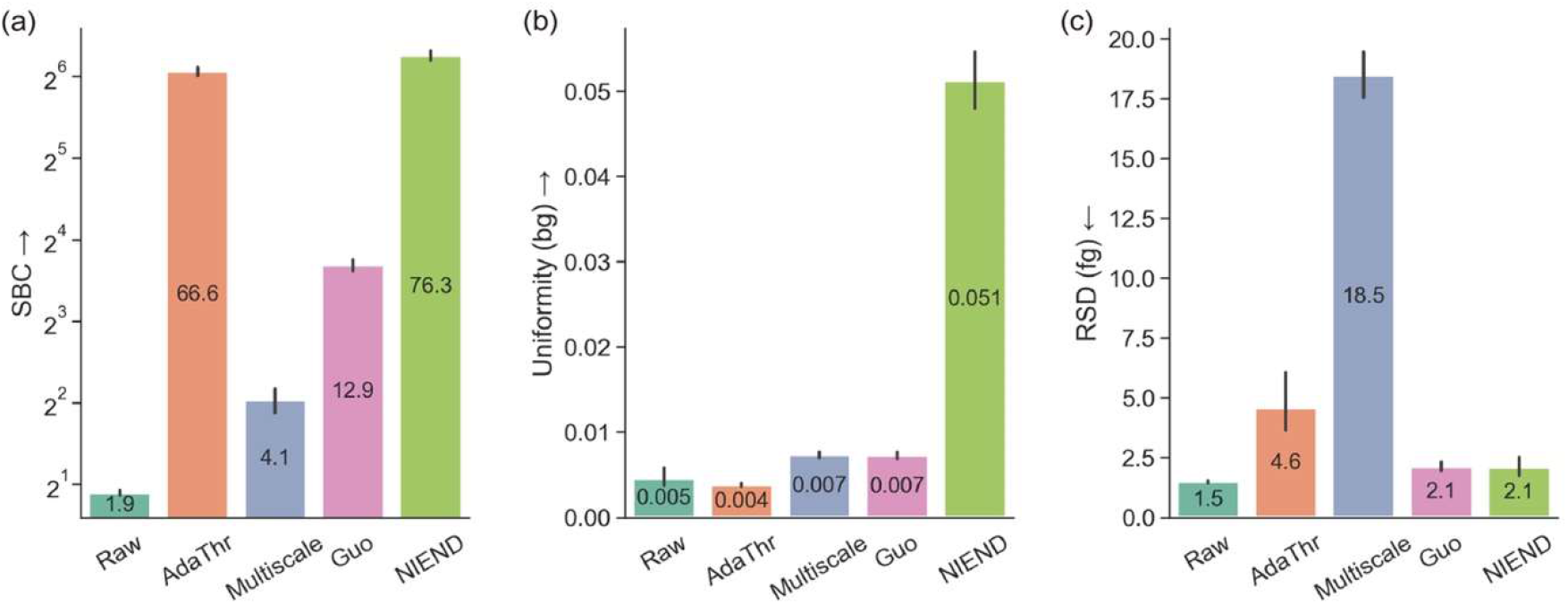
Quantitative comparison of enhanced image quality. (**a**) log-scaled signal-background contrast (SBC). (**b**) background uniformity (calculated on z-normalized intensities). (**c**) foreground relative standard deviation (RSD). The mean values are annotated onto the bars, and the error bars represent the 95% confidence interval for each mean. The direction of the arrow indicates better performance: an upward arrow means the higher the value, the better, and a downward arrow means the opposite.

### 3.3 NIEND improves automatic tracing

Automatic tracing on the processed images is benchmarked across the various image enhancing methods. In line with image quality results (**Figure 5**; **Figure 6**), adaptive thresholding could disrupt neurite signals near high-intensity regions, such as the soma, and thus may fail to reconstruct a large portion of arbors from the main stems (**Figure 7b,d,g**). multiscale enhancement results in greater disruption of neurite signals, leading to more significant arbor loss (the fourth column of **Figure 7**). Guo’s method yields a substantial improvement in recall, but some images exhibit over-tracing (**Figure 7e,g**). In contrast, NIEND outperforms in both avoiding over-tracing (**Figure 7d,e,g**) and retrieval of neurite structures (**Figure 7a,b,c**).

**Figure 7.**
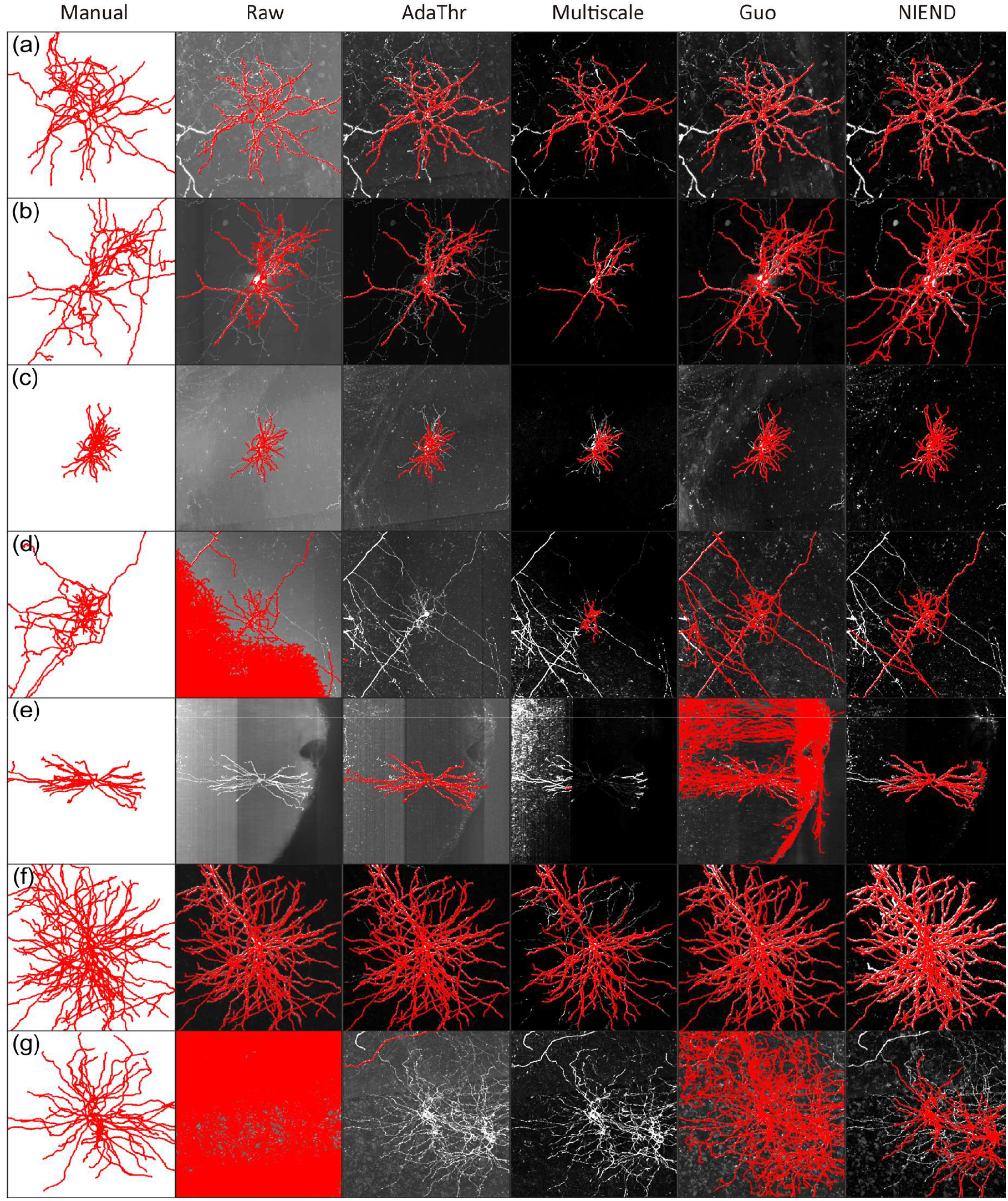
Comparison between NIEND and existing methods in automatic tracing. Each column represents one method, and each row corresponds to an image. The layout is analogous to that in **Figure 5**, except for the first column, which provides the manual annotations (gold standards). The reconstructed morphologies are overlaid onto the images and colored red. All images are z-score normalized and converted to unsigned 8-bit range (0 - 255).

We evaluated the precision, recall, and F1 of the tracing results on the 863 sparse neurons, and compared with existing methods (**Figure 8**). NIEND exhibits notably higher F1 scores than the raw image, adaptive thresholding, and multiscale enhancement, and holds a statistically significant advantage over the second-best Guo’s method (+4.4%). Adaptive thresholding and multiscale enhancement significantly outperform others in terms of precision, but their recalls are substantially smaller.. NIEND achieves a recall rate comparable to Guo’s method but offers superior precision (+4.3%), thus bringing a higher F1. We also assessed the reconstructions of NIEND-enhanced images using various topological metrics, confirming NIEND’s superior performance in reducing topological errors, including path fusion, path breaks, and loss of directionality (**Supplementary Figure S6**). We also performed the analysis on the BigNeuron dataset and using different tracing algorithms (**Supplementary Figure S8**). It is validated that NIEND can achieve comparable performance on other image modalities and the bonus is not specific to APP2. In summary, NIEND offers substantial improvement to automated tracing.

**Figure 8.**
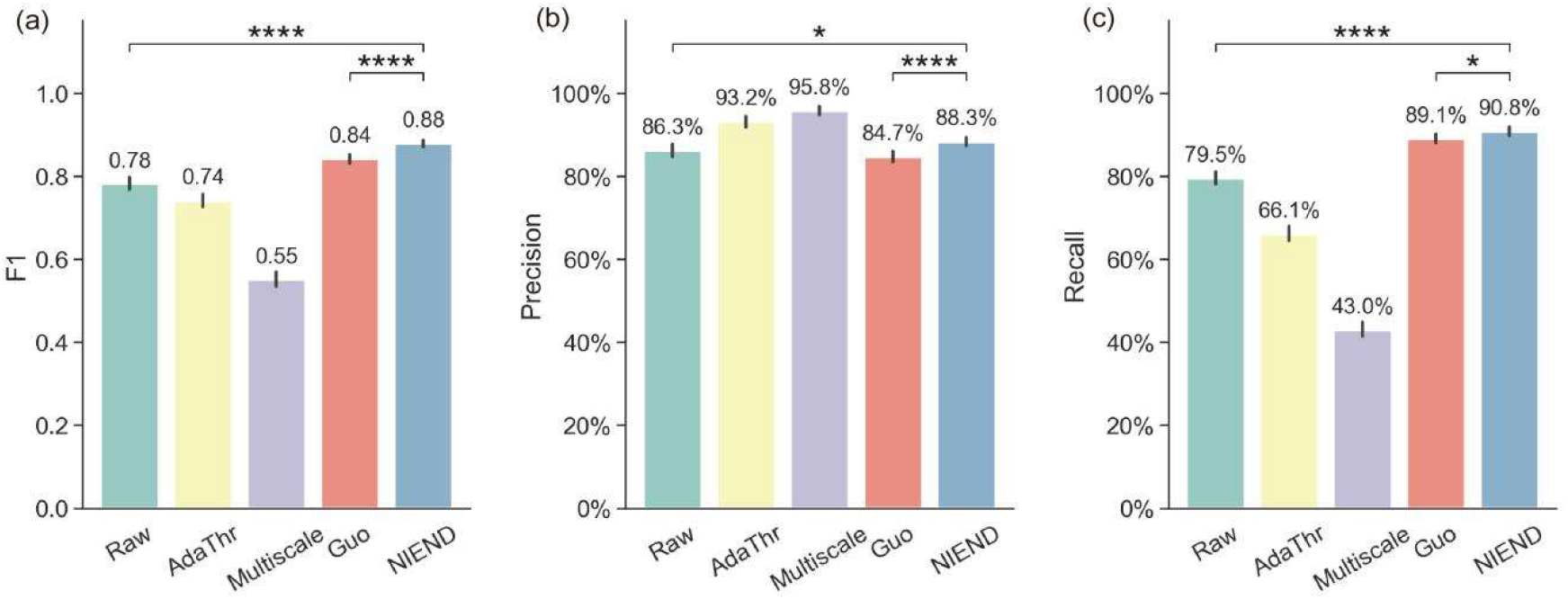
Quantitative evaluation of automatic traced morphologies. Bar plots of the F1 score (a), precision (b), and recall (c) for auto-traced morphologies of various enhancing methods. The error bar represent the 95% confidence interval for each method. Statistical significance between categories was tested (tow-sided T-test), and significance levels are indicated as follows: * for p < 0.05, ** for p < 0.001, *** for p < 0.001, and **** for p < 0.0001.

The comparison of image quality and tracing performance are furthered with R-L deconvolution (Supplementary Figure S9). Interestingly, it manifests more deterioration than enhancement from both aspects. We observed that R-L could produce extra artifacts around the neurite (**Supplementary Figure S9h**), leading to over-tracing and drop in precision. There could be many factors causing such failure, including the mismatch of PSF, the insufficiency of iteration, etc. Meanwhile, artifacts unrelated with PSF cannot be fully suppressed. Although improvement can still be made for the R-L deconvolution in our practice, the excessive amount of computation it demands is already dissuasive.

### 3.4 Ablation study

Ablation study is performed to assess the contributions from each step in the NIEND. The results are presented in **Supplementary Table 2**. Initially, the 16-bit images are substituted with 8-bit images. The tracing morphologies shows a considerable drop in recall (22%). This is not surprising as high-pass filtering (NIEND’s first step) can reveal more information from 16-bit images. The ablation of the diffusion filter displays minor drop in F1. It may be caused by the overlap of denoised targets between them. As illustrated in **Supplementary Figure S1b**, the diffusion filter can actually mitigate orthogonal artifacts to a certain degree. This overlap is further confirmed by the 4% decrease in recall when both steps are ablated. On the other hand, without the orthogonal filer, F1 even increases a bit (<0.001), indicating that its improvement and degradation are equally matched in our test samples and it yields little benefits in the presence of the diffusion filter. However, it is still adopted by NIEND for extra robustness. The intensity shifting exhibits the most significant impact on the tracing quality, with its ablation incurring a substantial 30% decrease in recall due to the loss of weak neurites. It is worth noting that adaptive thresholding suffers from a considerable decrease in recall (**Figure 8c**) for the same reason. The ablation of wavelet denoising shows only an insignificant drop. Similar to the orthogonal filter, wavelet denoising uniquely targets rare but intractable issues for other techniques. Another unintended consequence of wavelet denoising is the distortion of the neurite surface, to which some tracing algorithms are sensitive.

### 3.5 Performance advantage and high compression rate

Apart from the vast improvement in image quality and tracing accuracy, NIEND also facilitates a considerably low time and memory usage, which is, on average, 29.1s and 1GB per image for our tested cases (1024×1024×256). For images of a smaller size (256×256×256), they can be reduced to 1.6s and 64MB per image. The superior SBC and WIH (“uniformity” and “RSD”) offered by NIEND (**Figure 6**) are indicative of a high compression rate of the processed neuronal images. To compare the compression rate between different image enhancing methods, the images are compressed using the LZMA (Lempel–Ziv–Markov chain) algorithm in TIFF format (**Table 1)**. The initial size of each image (16-bit) used in this study is 512MB. Following each enhancing method, images are all standardized to 8-bit (256MB) and subsequently. Compression rates are computed as the compressed file size divided by the full file size. Of all the methods, multiscale enhancement achieves the smallest compression size (2.48MB), closely trailed by NIEND (2.49MB). Both methods achieve impressively low compression rates of 1.0%, surpassing all other methods. Considering the abnormally high signal loss of multiscale enhancement, NIEND emerges as the optimal choice for the efficient storage of petascale image data.

**Table 1.**
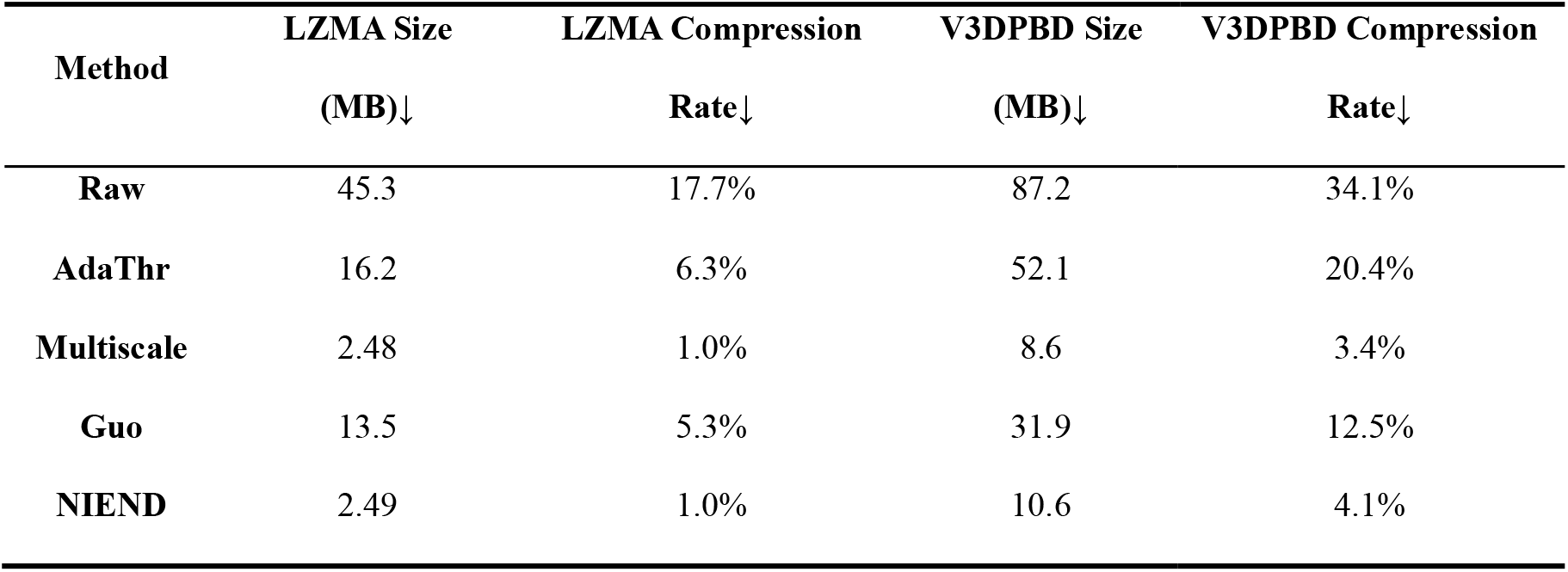
Average compression rate for 1891 256MB images using LZMA and V3DPBD compression of different methods.

We also evaluated the compression of V3DPBD image, a PackBits compression format for multi-dimensional images implemented by Vaa3D (Peng *et al*. 2014) and ImageJ (Schindelin *et al*. 2012). For images processed by NIEND, the file size can be reduced to approximately 4% (10.6MB) of the original sizes within seconds, a result that is only slightly less optimal than that achieved with the multiscale enhancer (8.6MB).

### 3.6 Whole-brain neuron tracing

Given its high efficiency and performance, NIEND can be easily integrated into the current pipelines of whole-brain single neuron tracing to boost the tracing performance. We showcased the tracing result of an exemplar single neuron using the APP2-based UltraTracer (Peng *et al*. 2017a) on a NIEND enhanced image (**Supplementary Figure S7**). Results show that the tracing of non-soma image blocks exhibit high accuracy in both noise removal and automatic tracing, resulting good single neuron tracing.

## 4 Conclusion and Discussion

In this work, we proposed a novel image enhancing pipeline, NIEND, for the enhancement of light-microscopic neuronal images. It effectively improved the image quality at a low time and memory cost, tackling various types of noise and artifacts resulting from technical flaws in the current sample preparation and imaging system. As a result, it succeeds in facilitating automatic tracing of higher performance and improving large-scale brain data curation.

Given the vast amount of imaging data of the mammalian nervous system, there is a significant need for either semi or fully automated pipelines for neuron morphology analysis. Numerous solutions have been proposed utilizing cutting-edge methods (Gao *et al*. 2022; Li *et al*. 2023), yet their outcomes are seriously limited in accuracy and efficiency, owing in part to the image quality issues. Our method approaches this dilemma as a cost-effective module adaptable. In comparison with sophisticated segmentation or other enhancing methods, NIEND is favorable for its high efficiency and low storage requirements, making it suitable for large-scale 3D biological images.

NIEND also turns the scale in favor of low-complexity tracing algorithms that have been limited to high-quality images. The low complexity typically relies on simplicity in resolving the image (e.g., the thresholding strategy of APP2) at the cost of robustness against noises and artifacts, which can be ameliorated with proper measures, e.g., the high-pass filtering of NIEND (**Supplementary Figure S5**). In this case, by applying APP2 thresholding on the sub-blocks of an image, the confidence of thresholding can be reflected by the standard deviation of thresholds. It is found that the standard deviation of thresholds drops after NIEND high-pass filtering, suggesting that the confidence is increased.

In terms of its novelty, NIEND achieves a good compromise between the model-based deconvolution and the frequency-based filtering, lowering the computational requirements while fortifying the specificity. The idea is inspired by Empirical Mode Decomposition (Huang *et al*. 1998), a common technique for non-stationary signal processing. In neuronal images, the non-stationary signal often emerges as irregular noises and artifacts, distorting the local statistical properties (e.g., the APP2 thresholding). As aforementioned, NIEND improves the APP2 thresholding confidence, by unifying the local statistical properties.

Although NIEND is theoretically specialized for fMOST images, its strong adaptivity allows it to be successfully applied to other modalities. Tailoring specialized solutions for other modalities by modifying the design of NIEND is also very simple because the source code of NIEND is light-weight. Overall, we hope NIEND can become an inspiration for more powerful image processing approaches in the future.

