## Supplementary Materials for "NIEND: Neuronal Image Enhancement through Noise Disentanglement"

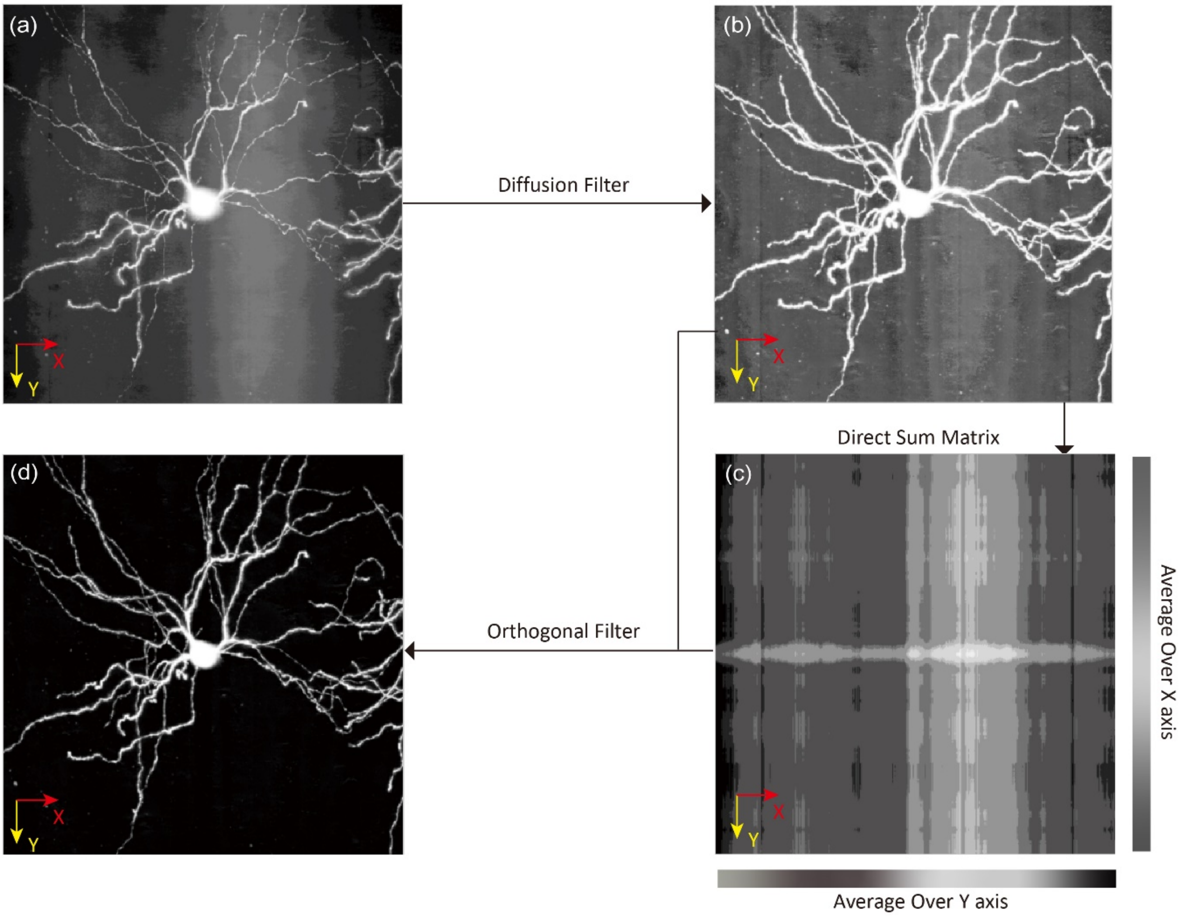


***Supplementary Figure S1.*** **Illustration of the orthogonal filter.** After diffusion filtering, an orthogonal filter is applied to address sinusoidal band-like artifacts. (**a**) A raw image with noises. (**b**) The image after diffusion filter. The diffusion filter substantially mitigates the artifact, yet the noise remains. (**c**) The maximal intensity projection (MIP) of the direct sum image. A direct sum image is the composition of the direct sums of all Z slices, each is constructed by the average intensities over the X-axis and that over the Y-axis for each slice. (**d**) The image after subtracting the direct sum image.

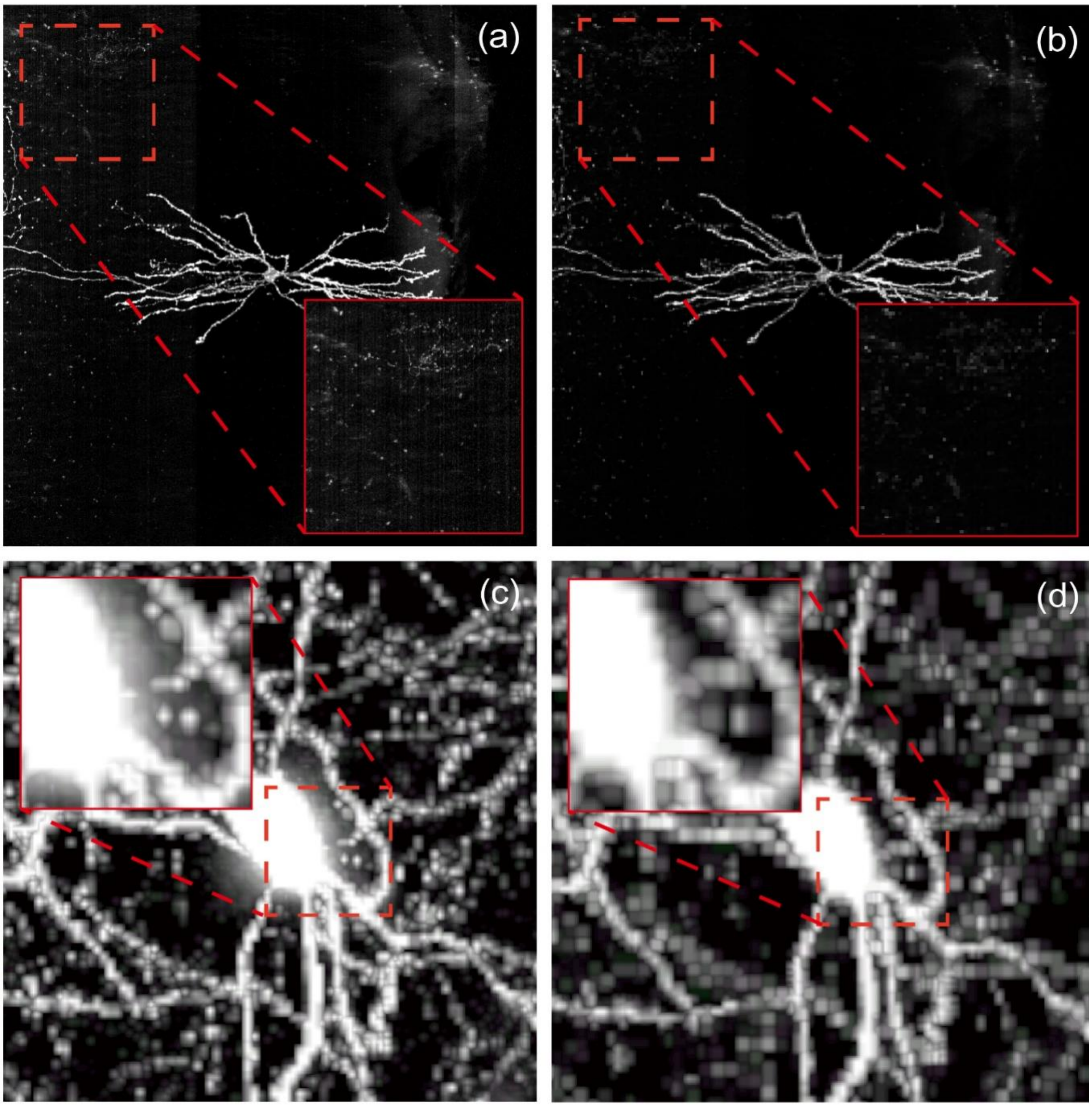


***Supplementary Figure S2.*** **Examples of wavelet denoising.** The wavelet denoising mitigates the glitches while preserving the integrity of neurites. (**a**) Elimination of the noises overlooked in previous stages. (**b**) Eradication of the remaining attenuation noises, and rectification of the discontinuous neurites.

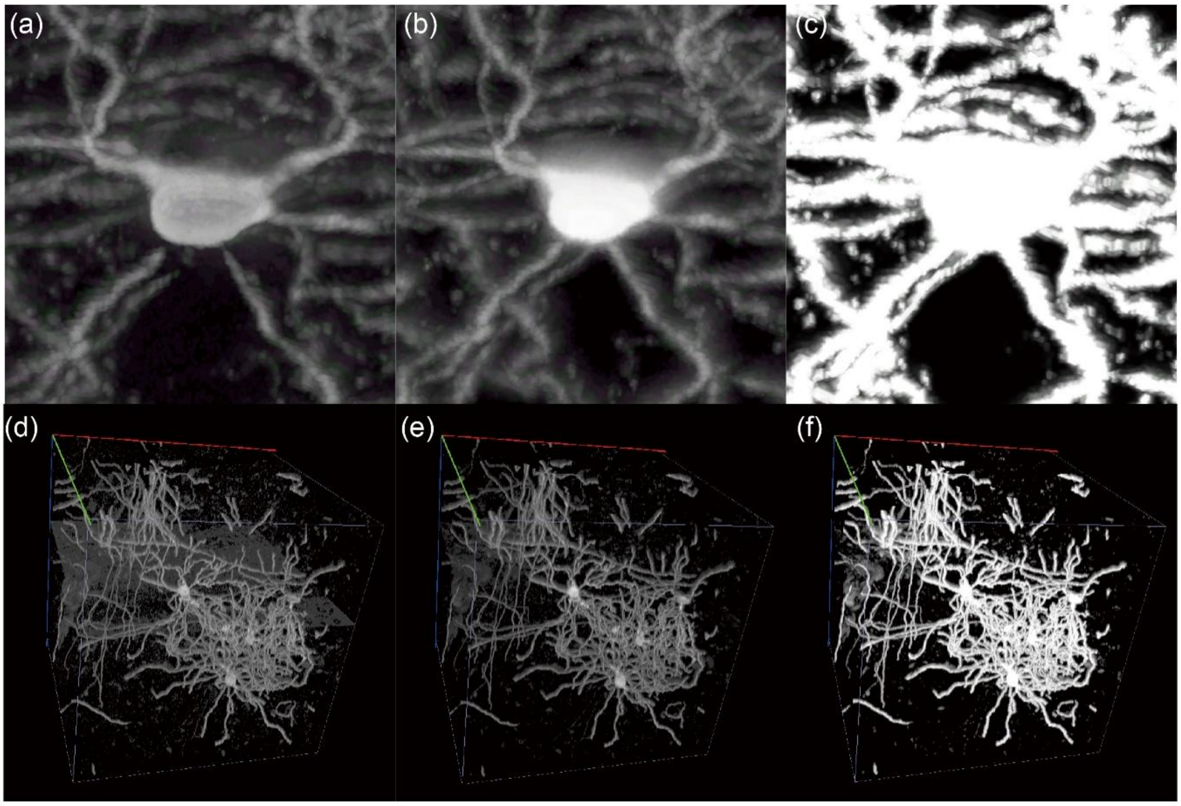


***Supplementary Figure S3.*** **NIEND outperforms adaptive thresholding in terms of high-pass filtering.** High-pass filtering is essential in image denoising. NIEND diffusion filter outperforms adaptive thresholding in denoising thanks to its improved adaptability to neuronal image artifacts. This figure showcases two scenarios where NIEND handles the artifacts by avoiding neurite disruption in denoising flare noises (**a–c**) and suppressing the horizontal boundary artifact (**d–f**). (**a**) Adaptive thresholding disrupts stems. (**b**) The diffusion filter attenuates the disruption. (**c**) The final result of NIEND shows the integrity of the neurites. (**d**) Adaptive thresholding does not address the planer artifact. (**e**) The diffusion filter in NIEND suppresses the artifact. (**f**) The final result of NIEND.

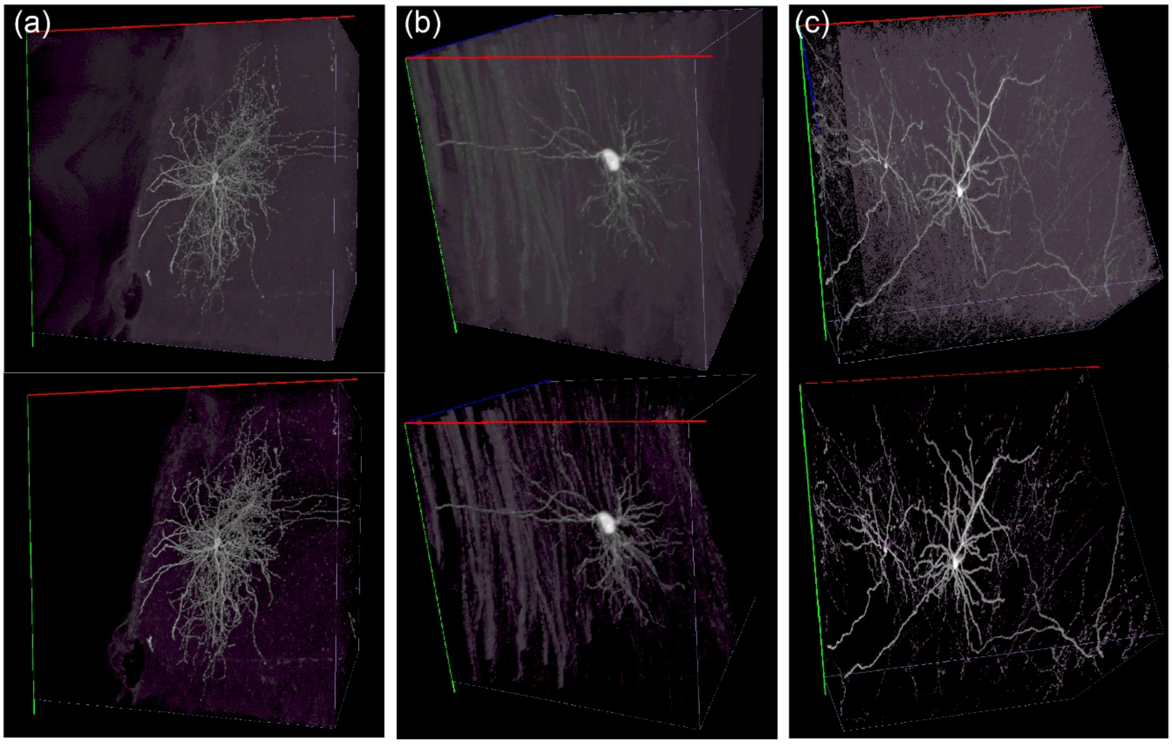


***Supplementary Figure S4.*** **Examples of images after the orthogonal filtering.** The orthogonal filter addresses large artifacts including but not limited to uneven illumination (shown in **Supplementary Figure S1**) and boundary artifacts introduced by image stitching or large tissues. The top row shows the input images and the bottom row shows the enhanced images. (**a**) An image with a large bulk of homogeneous noises. (**b**) An image with fibrous background noises. (**c**) An image with diffusive noises and clear boundaries.

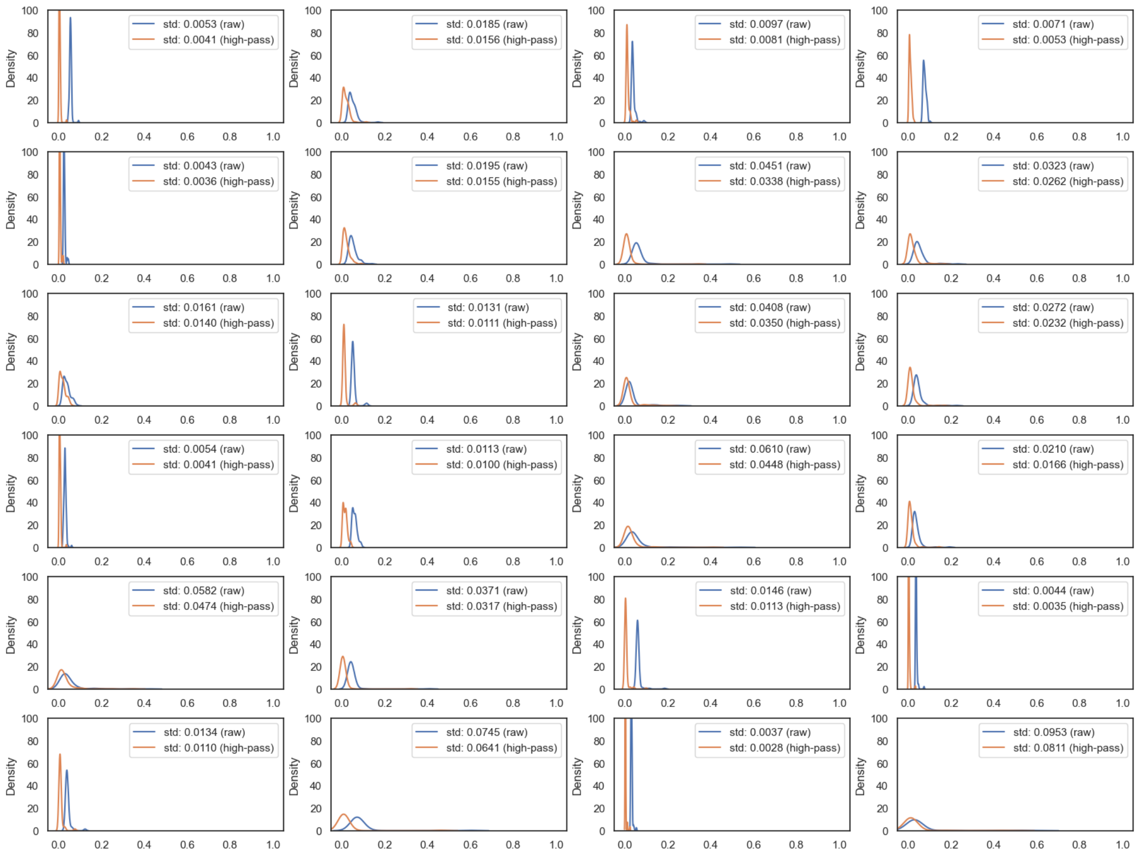


***Supplementary Figure S5*. Improvements in the local statistical property after NIEND high-pass filtering.** NIEND favors low-complexity tracing algorithms (e.g., APP2) by unifying the local statistical property, which is adopted in here as APP2 automatic thresholding, i.e., $\boldsymbol{Mean} + \mathbf{0.5} \cdot \boldsymbol{Std}$, where $\boldsymbol{Mean}$ and $\boldsymbol{Std}$ are the mean and standard deviation of an image region. To do this, 100 mini-blocks containing neurites with a size of $64 \times 64 \times 16$ are sampled and analyzed in 24 randomly chosen neuronal images. It is presented here their distributions before (blue) and after (orange) NIEND high-pass filtering, and the legends give the drops in the standard deviations of the thresholds. All values within each distribution are normalized to 0–1. The standard deviations in all of the 24 image blocks decrease after filtering, with an average reduction of 18%.

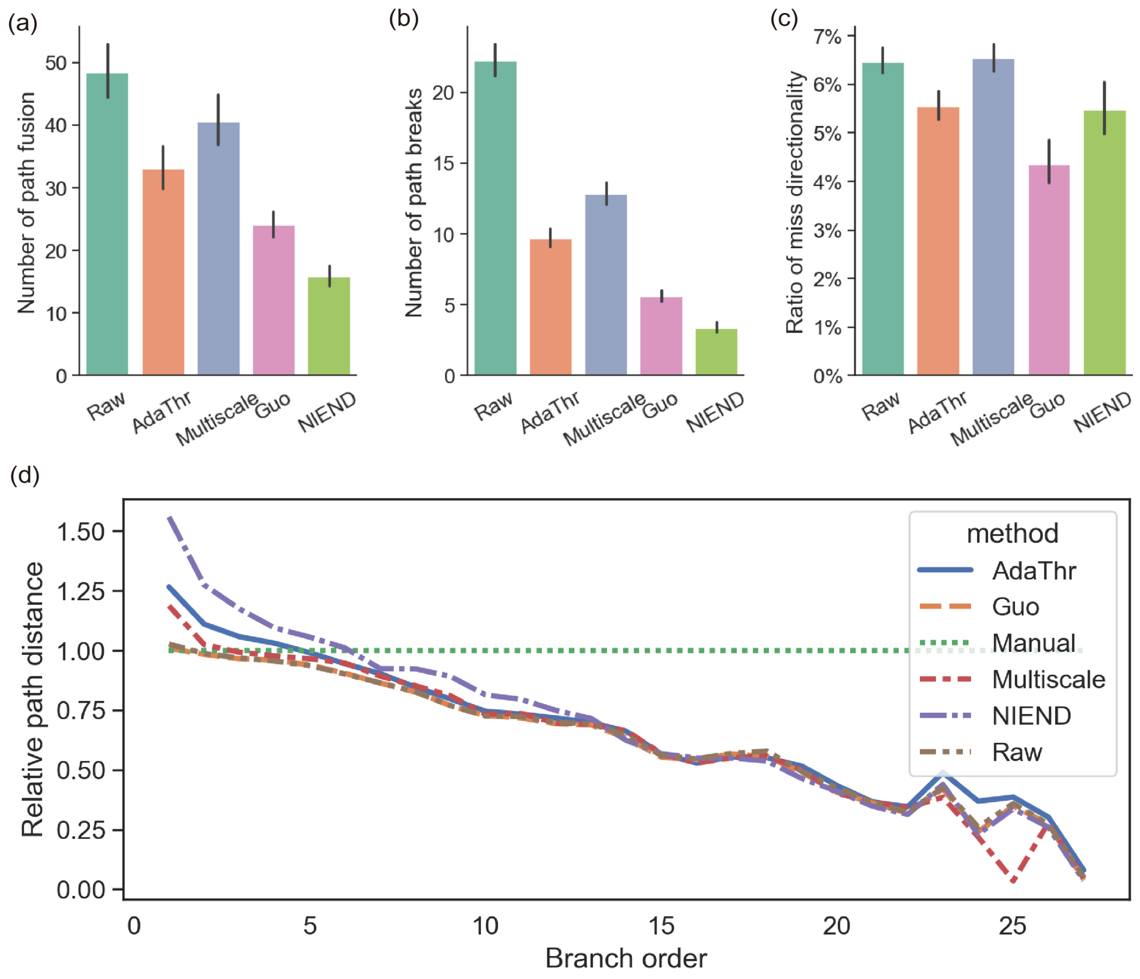


***Supplementary Figure S6.*** **Comparison of the topological measures of the automatically traced morphologies.** We compared different enhancing methods by quantifying four topological measures, including the number of path fusion (**a**), the number of path breaks (**b**), the ratio of miss directionality (**c**), and the path distance at each dendrite branch order relative to the gold standard (**d**). In each bar plot, the height of each bar represents the mean value, and the error bars represent the 95% confidence interval. The quantification is performed on each neuron and aggregated to represent the overall distribution for each method. The results reflect the increased robustness against topological errors of NIEND-processed images.

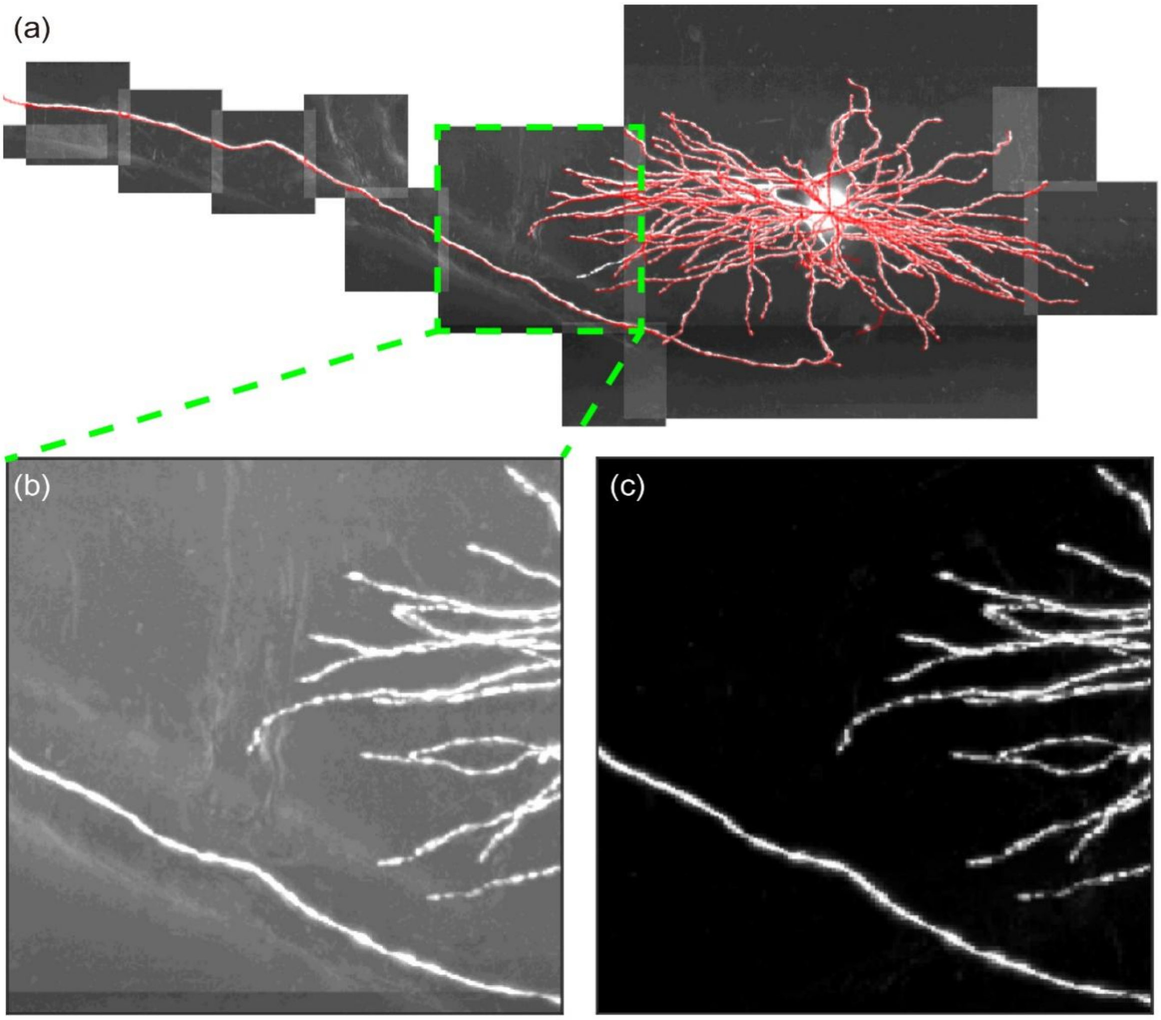


***Supplementary Figure S7.*** **NIEND-enhanced scalable tracing using UltraTracer.** (**a**) A single sparse neuron with a noisy background is processed with NIEND and progressively traced using UltraTracer (APP2-based). Crops are shown as raw images and z-normalized for better visualization. Zoom-in views of the exemplar blocks are shown in both its raw image form (**b**) and NIEND-enhanced form (**c**).

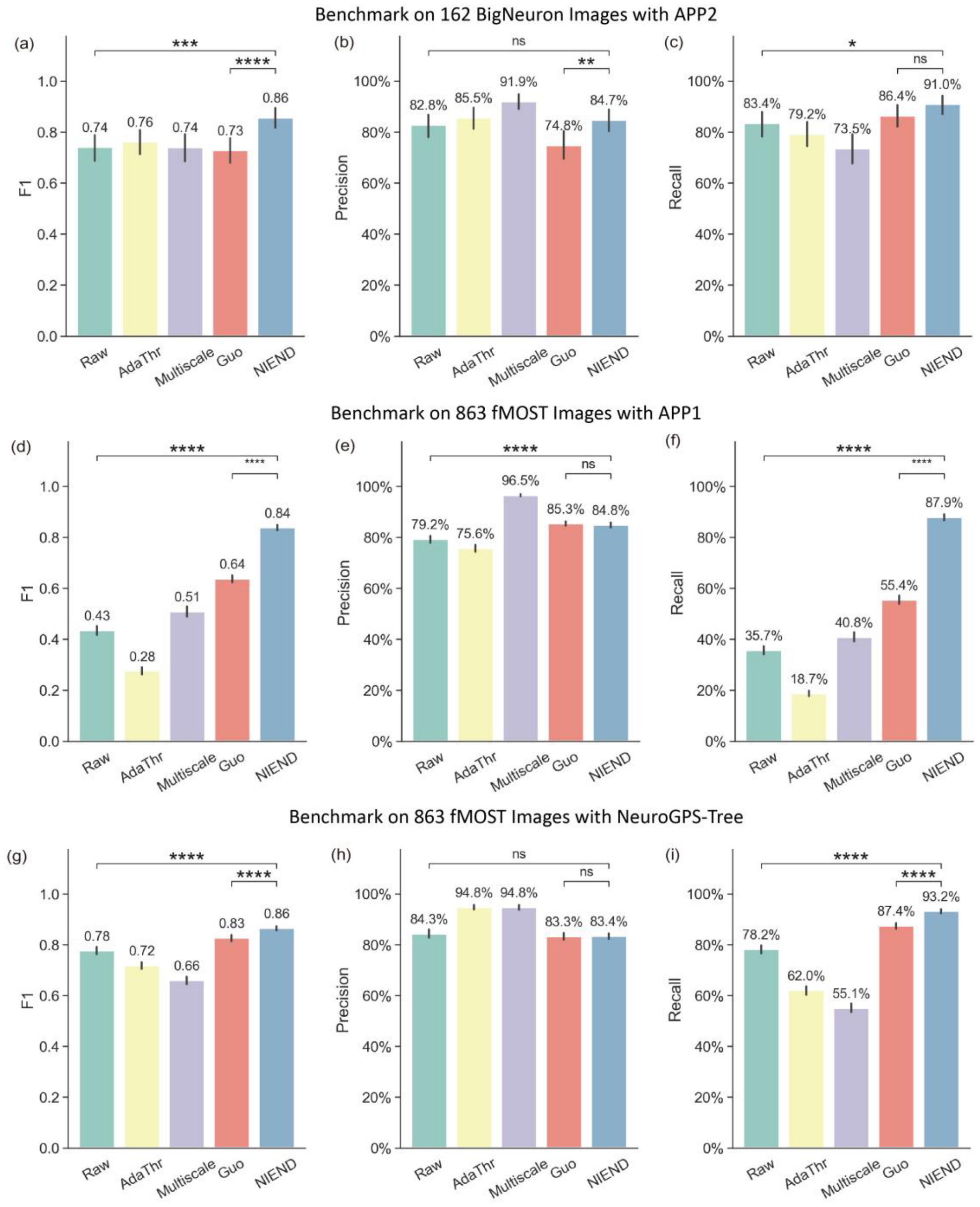


***Supplementary Figure S8.*** **Automatic tracing bench-testing.** In supplementary to the automatic tracing bench-testing shown in **Figure 8**, we also conducted bench-testing on the BigNeuron dataset and compared with other tracing algorithms such as APP1 and NeuroGPS-Tree. (**a**–**b**) Benchmark on the BigNeuron dataset highlights the superior performance of NIEND in image modalities other than fMOST. NIEND also leads in reinforcing APP1 (**d**–**f**) and NeuroGPS-Tree (**g**–**i**). In each component, the height of each bar represents the mean value for each method, and the error bars represent the 95% confidence interval. Statistical significance between categories was tested (two-sided T-test), and significance levels are indicated as follows: * for $p < 0.05$, ** for $p < 0.001$, *** for $p < 0.001$, and **** for $p < 0.0001$.

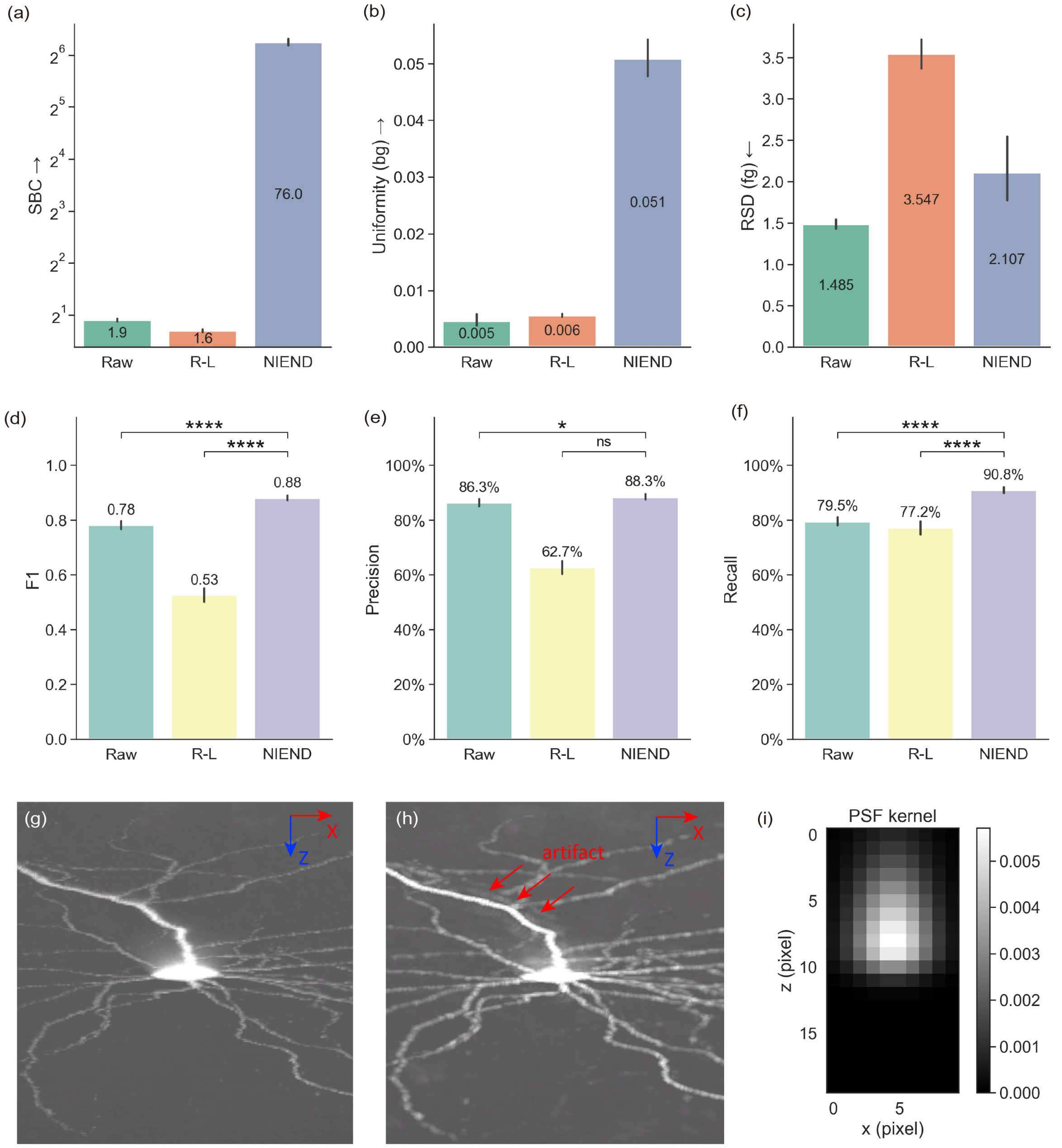


***Supplementary Figure S9.*** **Comparison with Richardson-Lucy deconvolution.** We compare the quality of the enhanced images processed with NIEND and Richardson-Lucy (R-L) deconvolution on 863 neurons. The first row displays the image quality metrics, and the second row shows the APP2 tracing accuracies. (**a**) NIEND achieves a significantly improved contrast, but R-L shows no improvement. (**b**) NIEND brings more improvement in background uniformity than R-L. (**c**) The loss of foreground uniformity is higher for R-L than NIEND. (**d**) In terms of F1, R-L worsens the tracing performance. (**e**) The precision under R-L is significantly lower than NIEND. (**f**) The recall under R-L is comparable to the raw image. These collectively demonstrate that NIEND can improve the image quality better than R-L and achieve better tracing results. The third row gives an example illustrating the performance of R-L. (**g**) An exemplar neuronal image. (**h**) R-L deconvolution of the input image. Although R-L enhances the neurite signal slightly, it introduces extra artifacts (marked by red arrows). (**i**) The side view of the PSF kernel (skewed Gaussian distribution, normalized by the sum of the kernel) applied on all the neuronal images.

### Supplementary Methods

#### Preparing the BigNeuron dataset

The BigNeuron images are downloaded from http://web.bii.a-star.edu.sg/bigneuron/gold166.zip (containing 164 fluorescent images). Then, images of incompatible sizes (intractable by benchmarking image processing algorithms implemented as Vaa3D plugins) are excluded, with 162 images left. Afterwards, the dataset undergoes the same processing and analyzing pipeline as the 863 neuronal images (**Methods**).

#### Auto-tracing with APP1, APP2 and NeuroGPS-Tree

APP1, APP2 and NeuroGPS-Tree are performed on each of the 863 neuronal images with the same parameter settings. Both of APP1 and APP2 use automatic thresholding ($\boldsymbol{Mean + 0.5 \cdot Std}$, where $\boldsymbol{Mean}$ and $\boldsymbol{Std}$ are the mean and standard deviation of the image intensities) and automatic downsampling is turned off (with this on, large image blocks will be resized as 256×256×256), with other parameters left as default. Manual annotated soma positions are utilized as the initial points for the automatic tracing.

For NeuroGPS-Tree, the resolution is set as 0.25/0.25/1 for *x*/*y*/*z* axes. Additionally, the binarization threshold is set as 1; the trace value is set as 6; and the enhance value is set as 0.

#### Deconvolving using the Richardson-Lucy method

The Richardson-Lucy deconvolution (R-L) is performed by first modeling the point spread function (PSF) and then conducting the iterative deconvolution. Due to the inaccessibility to experimental settings of the microscope of each brain, we use an axially skewed 3D Gaussian distribution to approximate the PSF. It is generated by multiplying a lateral 2D Gaussian distribution and an axial 1D skewed Gaussian distribution. The lateral standard deviation is set as 2. The skewness of the axial distribution is set as 5. The scale of the axial distribution is set as 5. The skewed Gaussian distribution is defined as $\boldsymbol{2 \cdot NormPDF(x) \cdot NormCDF(Skewness \cdot x)/scale}$, where $\boldsymbol{NormPDF}$ stands for the standard Gaussian probability distribution function, $\boldsymbol{NormCDF}$ stands for the standard Gaussian cumulative distribution function, $\boldsymbol{Skewness}$ determine the skewness and $\boldsymbol{Scale}$ scales the distribution. The window size of the PSF is set as 10 pixels in X- and Y-axis and 20 pixels in the Z-axis. Using the modeled PSF, the deconvolution is performed by 30 iterations.

#### Breaking down the topologies of reconstructions

To quantify the topological accuracy, 4 metrics are computed on each automatic traced morphologies of the 863 neuronal images and compared with human-annotated gold standard. They are defined as follows:

**The number of path fusion**. Path fusions are incorrect connections between neurites, leading to a different path to soma than the gold standard. For each critical node, we approximate the identification of path fusion by finding its nearest node on the reconstructed neuron and compare their path distances to soma. If the difference between the path distances exceeds 10 pixels and the ratio between them exceeds 1.2, the critical node will be marked as a path fusion.

**The number of path breaks**. Path breaks are breaks along a neurite that should be continuous. The quantification of path breaks is performed by assessing the continuity of the counterpart of each branch in the gold standard. A branch is defined as the path between two adjacent critical nodes (branch nodes and tips). The counterpart is located by matching the nearest nodes for the two critical nodes. When the path distance between the two matched nodes is 10 pixels or 1.2 times larger than that in the gold standard, it is counted as a path break.

**The ratio of miss directionality**. When a topological error occurs, the traced neurite can go along a direction opposite to that in the gold standard. The quantification of such miss directionality is performed by assessing all the node directions on the gold standard. For each node on the gold standard, the nearest node can be found on the tracing result. The direction of a node is defined as the vector from its parent node to itself. If the angle between the directions of the 2 matched nodes exceeds 90°, it will be marked as reversed and the length of the vector (in gold standard) will be summed up. Finally, a total reversed path length will be calculated.

**Path distance to soma at each branch order (dendrite only)**. Since the branching patterns are different among the traced morphologies, the branch orders can be inconsistent. Therefore, we only take the branch nodes from the gold standard and find their nearest nodes in each tracing result to determine their path distances.

For all the above analyses, the distance for any two matching nodes must be within 5 pixels.

### Tracing single neurons with UltraTracer

To test the capacity of NIEND enhancing in single neuron tracing, we extracted the image of a manually annotated neuron from the whole brain image and traced using UltraTracer. First, it is cropped as a down-sampled block of 2048×2048×256 voxels, with its resolution as 0.40μm/pixel in the lateral directions and 2μm/pixel in the axial direction. Then, the image block is processed with NIEND and converted to the TeraFly format using Vaa3D TeraConverter. Finally, UltraTracer (APP2-based) is performed with the initial block size of 512×512×128. UltraTracer automatically adaptively detects and crops new image blocks and traces based on previous tracing results until all candidate blocks are reconstructed.

***Supplementary Table S1***. Parameter configurations of the diffusion filter and the intensity shifting

| **Lateral Resolution (µm/pixel)** | **s** | **Percentile Threshold (foreground)** |
|---|---|---|
| 0.2 | 15 | 1.5% |
| 0.24 | 12 | 1% |
| 0.35 | 8 | 0.5% |

***Supplementary Table S2***. Average F1, precision, and recall for the ablation of each NIEND step

| Method | Average F1↑ | Average Precision↑ | Average Recall↑ |
|---|---|---|---|
| **full** | 0.8792 | 88.33% | 90.84% |
| **8-bit image** | 0.8348 | 93.60% | 78.76% |
| **w/o diffusion** | 0.8639 | 88.79% | 87.78% |
| **w/o orthogonal** | 0.8797 | 88.39% | 90.81% |
| **w/o high-pass** | 0.8540 | 88.91% | 86.05% |
| **w/o intensity shifting** | 0.7701 | 95.14% | 69.16% |
| **w/o wavelet** | 0.8788 | 88.70% | 90.26% |
